# Selective over-synthesis and rapid turnover of mitochondrial protein components of respiratory complexes

**DOI:** 10.1101/832287

**Authors:** Daniel F. Bogenhagen

## Abstract

Mammalian mitochondria assemble four complexes of the respiratory chain (RCI, III, IV and V) by combining 13 polypeptides synthesized within mitochondria on mitochondrial ribosomes (mitoribosomes) with over 70 polypeptides encoded in nuclear DNA, translated on cytoplasmic ribosomes and imported into mitochondria. We report that pulse-chase SILAC can also serve as a valuable approach to study RC assembly as it reveals considerable differences in the rates and efficiency of assembly of different complexes. While assembly of RCV, ATPase, was rapid with little excess synthesis of subunits, RCI, NADH dehydrogenase, assembly was far less efficient with dramatic over-synthesis of numerous proteins, particularly in the matrix exposed N- and Q-Domains. Subunits that do not engage in assembly are generally degraded within three hours. Differential assembly kinetics were also observed for individual complexes immunoprecipitated with complex-specific antibodies. Immunoprecipitation with an antibody that recognizes the ND1 subunit of RCI co-precipitated a number of proteins implicated in FeS cluster assembly as well as newly-synthesized UQCRFS1, the Rieske FeS protein in RCIII, reflecting a degree of coordination of RCI and RCIII assembly.

---

Mitochondria maintain a complete genetic system dedicated to the synthesis of a small number of components of the respiratory chain within the organelle. The 16.5 kB mammalian mtDNA genome encodes 13 of the ∼80 subunits of the oxidative phosphorylation system. In a unique gene organization, the mRNAs for these proteins are co-transcribed with the rRNAs and tRNAs required for their translation. Thus, mitochondrial biogenesis depends on two parallel assembly lines to build both the respiratory chain and the mitoribosomes.

Mitoribosomes are constructed by orderly addition of 80 nuclear-encoded proteins to the rRNA scaffolds in a process that is nearly as complex as assembly of the entire respiratory chain. During the course of experiments to investigate the kinetics of mitoribosome assembly in HeLa cells, we used stable isotope pulse labeling in cell culture (pulse-SILAC) to study two aspects of the assembly process. First, we showed that a subset of newly-synthesized mitoribosomal proteins (MRPs) associated selectively with the mtDNA nucleoid, consistent with their binding to nascent rRNA transcripts (1). We then used SILAC in a pulse-chase labeling format to generate a coarse-grained model for assembly of the mitoribosome (2). One of the principles of this approach is that when a multi-subunit structure is analyzed after pulse-labeling the last proteins added to complete its construction have the highest content of pulse label. This method has the greatest resolution for assembly processes that require a time interval comparable to or longer than the pulse labeling interval. Using this approach, we were able to distinguish early, intermediate and late-binding proteins in mitoribosomes according to their labeling kinetics. We found that mitoribosome assembly requires a time interval of 2-3 hr. By comparing the pulse-label content of individual proteins in the total mitochondrial pool as well as in the mitoribosome, we were able to show that many MRPs are synthesized and imported into mitochondria in considerable excess. This is not commonly seen for proteins that are not part of large complexes (3), although different proteins can have different half-lives. The excess MRPs are turned over if they are not successfully assembled in a few hours. We interpreted this as a mechanism to ensure a sufficient supply of parts for the complex process of mitoribosome assembly. This strategy has obvious disadvantages: It is inefficient, since many protein molecules are effectively wasted, and it requires a robust selective apparatus to degrade the excess proteins.

We have now expanded this approach to ask whether this is a common feature of other mitochondrial macromolecular machines such as respiratory complexes (RC). Numerous previous studies of RC assembly have been valuable in establishing many aspects of the orderly mechanism of assembly for these complexes. However, many of these studies have used radioisotope tracers and/or have perturbed assembly by treatment for several hours with translation inhibitors cycloheximide or chloramphenicol in order to monitor the kinetics of assembly (4, 5). We undertook to determine whether SILAC labeling would provide a valuable supplement or alternative to these approaches.

Pulse labeling with stable isotopes of essential amino acids can distinguish newly-synthesized proteins from pre-existing copies. We grew HeLa cells in “light” medium containing ^12^C-lysine and arginine and switched to medium with lysine and arginine containing six ^13^C residues in each to initiate pulse labeling, continued growth for defined time periods and isolated mitochondria and/or respiratory chain complexes. Labeling with stable isotopes enables mass spectrometric detection of the heavy:light or H:L ratio of a large fraction of the mitochondrial proteome as an index of the synthesis rate for a far greater number of proteins than in a radioactive labeling experiment. When labeling is done on non-synchronized exponentially-growing cells, we can make the steady-state assumption that total cell protein should double during each cell generation and calculate the expected H:L ratio at various labeling times (2). We found that some RC, such as F_1_/F_o_ ATPase (Complex V) are assembled efficiently with little subunit wastage, while others include some subunits that are imported in considerable excess, as reflected by elevated H:L ratios. This apparent over-synthesis of proteins was particularly dramatic for some subunits of RCI (NADH dehydrogenase) with an effective wastage of several times as many protein copies as we observed during mitoribosome assembly. We suggest that pulse-chase SILAC labeling can provide a useful additional method to study complex assembly and protein degradation rates as well as the alteration of these dynamic processes in human mitochondrial diseases.

## Results and Discussion

### Rates of synthesis and turnover of mitochondrial proteins

We used replicate HeLa cell cultures to perform SILAC pulse-labeling experiments for 3, 4, 6 or 12 hr and isolated total mitochondrial proteins using differential centrifugation and sucrose gradient sedimentation. The proteins were fragmented with trypsin and peptides were analyzed by LC-MS/MS to determine the relative quantity of labeled (H) and unlabeled (L) peptides. The biochemical preparations and data handling procedures used to generate customized reports of protein H:L ratios were as described (2). Table S1 reports labeling data on several hundred proteins categorized as mitochondrial proteins in the Mitocarta database (6), although the coverage (number of peptides observed) of many less-abundant proteins is often low or inconsistent. We selected 19 abundant proteins that were not part of large multi-protein complexes as internal “standards” that showed similar labeling kinetics (Fig 1A). In general the rate of accumulation of newly synthesized protein slightly exceeds that predicted by our mathematical model (dashed blue line in Fig 1A), which calculates expected H:L ratios based on the cell replication rate assuming that pre-existing light proteins are relatively stable during our experiments as described (2). This heuristic assumption is, of course, incorrect due to protein turnover, which is a variable but minor factor for most proteins in short term labeling experiments. These standard proteins were labeled at a rate 18 to 50% higher than predicted by the model, which most likely reflects the excess synthesis and mitochondrial import of proteins required to replace copies lost to turnover. In comparison with some proteins discussed below, the standard proteins exhibited a tight variation with a standard deviation of H:L values less than or equal to +/-23% of the mean at each time point. In the subsequent analysis of respiratory complex proteins, we consider that proteins with an H:L ratio more than 2 standard deviations greater than the mean for these standard proteins can be considered over-synthesized. None of the 19 standard proteins exceeded this threshold.

**Figure 1.**
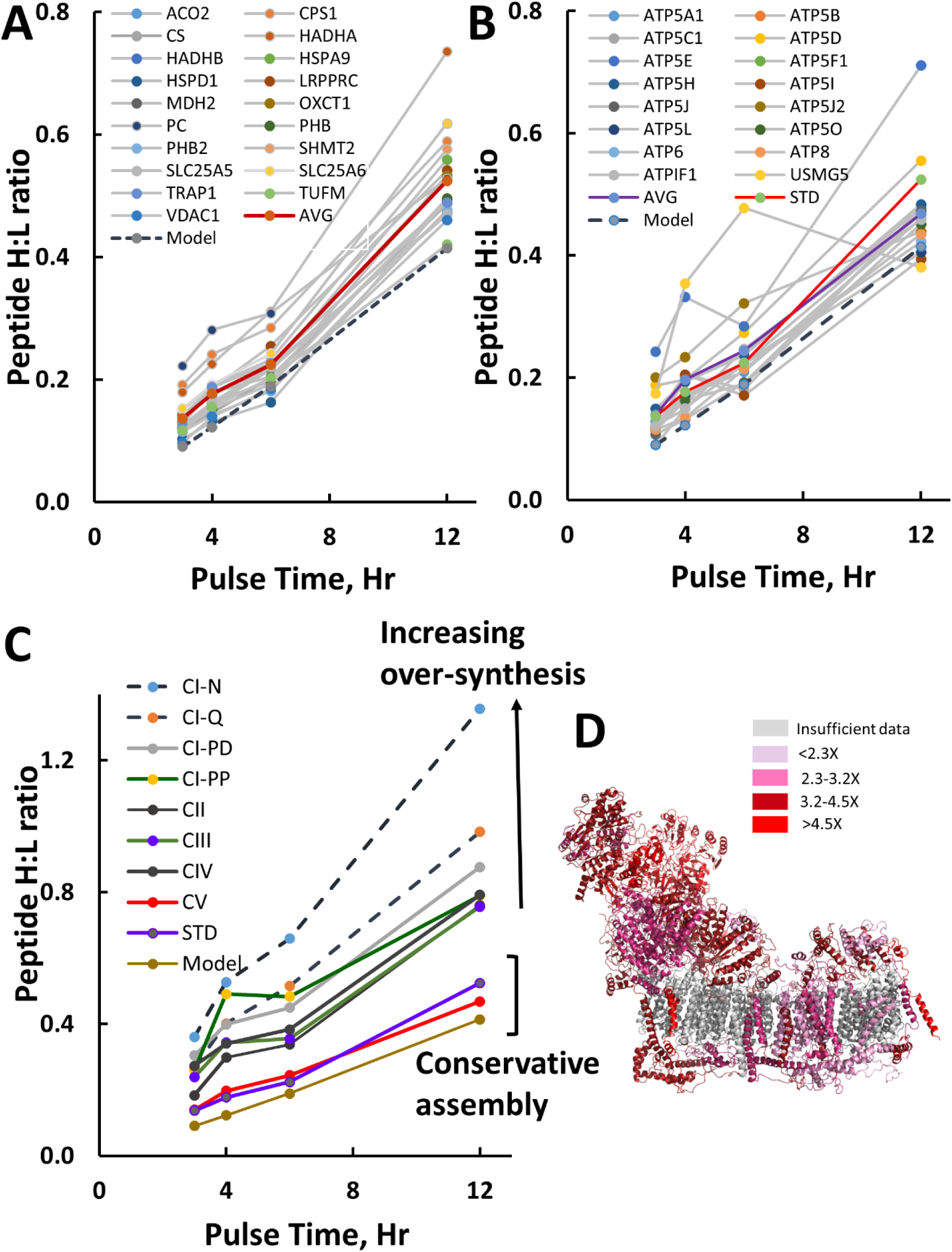
Kinetics of synthesis and import of mitochondrial RC proteins. Total HeLa cell mitochondrial proteins were isolated after SILAC pulses of 3, 4, 6 or 12 h and analyzed by fragmentation and LC-MS/MS. Average peptide H:L ratios are plotted as a function of labeling time. **A.** Representative standard proteins exhibit similar rates of accumulation of newly-synthesized protein with some variability surrounding an average shown by the red line. The dashed line shows the expected rate of accumulation of newly-synthesized proteins assuming exponential growth with a generation time of 24 h and negligible protein turnover. **B.** Subunits of Complex V, F_1_F_o_ATPase, on average (purple line) show synthesis kinetics similar to the standard proteins (red line). **C.** Comparison of the synthesis rates of subunits of all RC shows that RCIII and RCIV proteins are synthesized and imported more rapidly than RCV subunits, which are synthesized with minimal excess, and that RCI subunits exhibit even higher rates of over-synthesis, particularly for the matrix-exposed N and Q domains. Note that NDUFS6 is not included in this analysis since it is synthesized and incorporated at even higher rates, over twice that of the average RCI subunit. Higher rates of accumulation reflect progressively greater over-synthesis. **D.** Ribbon model of RCI based on RSCB PDB model 5LDW with individual subunits colorized according to their extent of over-synthesis in mitochondria from cells pulse-labeled for 3- and 4- h relative to the H:L ratios predicted by the model. Summary H:L ratio data are from Table S1. Molecular models were generated using Pymol.

We analyzed the protein accumulation kinetics observed for the 16 members of RCV, F_1_F_o_ATPase, including the two mitochondrially-synthesized components, ATP6 and ATP8 (Fig. 1B). The H:L ratios of these proteins as a function of labeling time revealed that they were synthesized and imported at rates very similar to the standard proteins. Although the data suggest slight over-synthesis of ATP5E and USMG5 (ATP5MD/DAPIT), the average H:L ratios for Complex V proteins were very close to those of the standards (Fig. 1B purple and red curves). We conclude that RCV proteins are synthesized and imported at a pace appropriate to keep up with cell proliferation with minimal excess synthesis or protein wastage.

Applying this same analysis to the other respiratory complexes showed that the rates of synthesis of many other RC subunits appreciably exceeded those of RCV subunits. We obtained sufficient peptides to analyze the rates of synthesis of 34 of the 44 RCI (NADH dehydrogenase) structural subunits and 15 assembly factors. We also had adequate peptide coverage to evaluate 27 of the 35 subunits of RC II, III and IV. Setting a threshold for accumulation rate two standard deviations above that for the standard proteins, we found that 34 subunits of respiratory complexes exceeded this threshold, including most subunits of RCI (Table 1). As we suggested in the case of mitoribosome assembly (2), the high rate of synthesis and import of individual subunits may serve to reduce delays in complex assembly that might occur is a particular polypeptide were not available when required. Complex I proteins accumulated at the highest rate, particularly those residing in the matrix-exposed N domain (Fig. 1C), with an average accumulation rate 2.7 times higher than RCV. Indeed, the only component of the hydrophilic arm of RCI that did not accumulate at an accelerated rate was NDUFAB1 which plays additional roles as the acyl carrier protein and as a component of the FeS cluster assembly complex (7). Fig. 1C omits NDUFS6 which exhibited an exceptionally high synthesis rate about twice as great as the other N-domain proteins (Table S1). This rate is sufficient to replace 75% of preexisting NDUFS6 in only 12 hr. Mammalian RCI is known to contain 44 subunits, far more than a typical bacterial NADH dehydrogenase, with the additional or supernumerary subunits often located surrounding a core of subunits closely related to their prokaryotic counterparts (8, 9). Mapping the rapidly accumulated RCI subunits within the structure of the complex (Fig. 1D) revealed that some of the supernumerary subunits in the membrane domain also accumulated more rapidly than the core subunits. RCII, III and IV as well as the membrane domains of RCI accumulated at intermediate rates, more rapidly than RCV, but far slower than the hydrophilic matrix-exposed N and Q domains of RCI. One interesting exception is NDUFA4 which is one of the most highly over-synthesized proteins (Fig. S1). Despite its name, this is part of RCIV (10) and a recent structural study suggested that it would be positioned in RCIV at a location that would block formation of RCIV2 dimers (11). It may be that maintenance of high levels of NDUFA4 through continual over-synthesis is necessary to occupy this site.

**Table 1.**
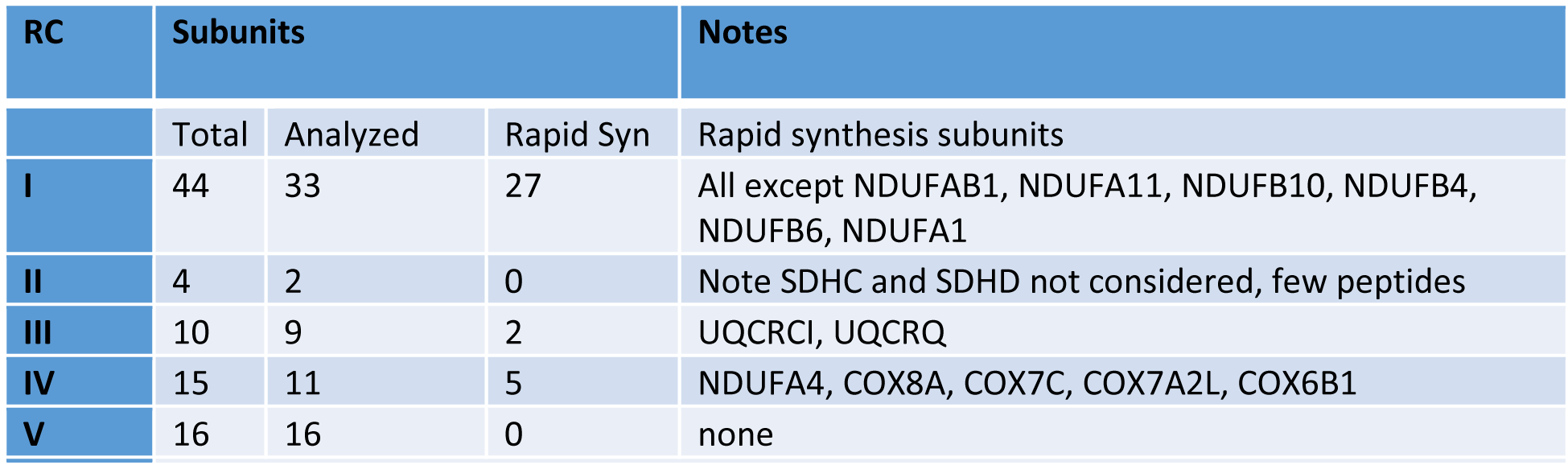
RC subunits rapidly synthesized and imported into mitochondria.

Since respiratory complex assembly involves the action of a large number of accessory factors, we asked whether these factors are generally accumulated with the same kinetics as the structural subunits. Because excessive synthesis is most apparent with the shorter labeling times of 3 or 4 hr, we compared the H:L ratios of the respiratory chain assembly factors with those of structural subunits at these labeling times (Fig. S1A). We found that assembly factors are not generally over-synthesized to the same extent as structural subunits of complexes. Evidently, since accessory factors are not consumed by assembly processes there is no advantage to their over-synthesis.

A high rate of increase in the peptide H:L ratio for a protein may indicate a high rate of turnover, either of the entire complex or of individual protein components. This is particularly relevant in cases where an individual protein might not be stably bound within a complex, but might be readily exchangeable, as has been noted in radioactive labeling experiments (5). To differentiate between these two possibilities we conducted pulse-chase SILAC experiments in which we labeled with ^13^C-lysine and arginine for 4 hr and chased with unlabeled medium for 3, 7 or 10 hr (Table S2). A 4-hr labeling was selected as a relatively short time period that would provide sufficient ^13^C incorporation to permit proteins to be monitored adequately. We performed some chase experiments following a 3-hr labeling interval (data not shown) that were generally consistent with the results obtained with 4-hr labeling, but were limited by the low initial H:L ratios of most proteins due to the short labeling interval. Fig 2A shows that the H:L ratios of the standard proteins declined during this chase interval as expected due to continued synthesis of unlabeled proteins during the chase. Protein turnover may also contribute to the decrease in H:L ratio during the chase, particularly for proteins like PC, CPS1 and HADHA that show a steeper decline in their H:L ratios during the first three hours of the chase in Fig. 2A. Fig. 2B showed that most RCV proteins behaved similarly, although USMG5 is characterized by higher than expected H:L ratios after the pulse and exhibited more rapid turnover, as noted in a recent comprehensive study of RCV assembly (12). This hallmark of turnover is exhibited even more dramatically by subunits of the other respiratory complexes. Fig. 2C shows that the average H:L ratios of RCI to IV subunits were higher than that for RCV immediately after the pulse. Note that the error bars are large immediately after the pulse, reflecting great variation in H:L ratios among individual proteins. This is shown for individual proteins in Table S2. The initially elevated H:L ratios generally converged to a lower, more consistent average during 3 to 10 hr of chase, with most of the decrease evident in the first 3 hr (Fig. 2C). This is particularly clear for the RCI N- and Q-domain proteins since many of these have high rates of synthesis. Fig. S1B,C show that the H:L ratios for these over-synthesized proteins quickly decline during the first 3 hr of chase. In general, individual proteins that rapidly achieve high H:L ratios in a 3- or 4-hr pulse are synthesized and imported at rates in excess of those required to maintain steady-state mitochondrial protein composition and are metabolically unstable. Some examples of other very rapidly synthesized proteins are shown in Fig S1D. It has previously been found that ALAS1, the rate-limiting enzyme in heme biosynthesis, is rapidly turned over in a heme-dependent manner (13). We are not aware of previous reports on the turnover of other proteins shown in Fig. S1B. The high rates of synthesis of NDUFS6 and the RCIV protein NDUFA4 have been noted above. The high rates of synthesis of COQ10B and TRUB2 have not previously been observed, to our knowledge. We conclude that newly synthesized imported copies of structural subunits of respiratory complexes are subject to biphasic turnover kinetics. The imported RC subunits may be either built into complexes or sub-modules, after which they are subject to a slow rate of turnover, or the excess unassembled copies are commonly degraded within about 3 hr.

**Figure 2.**
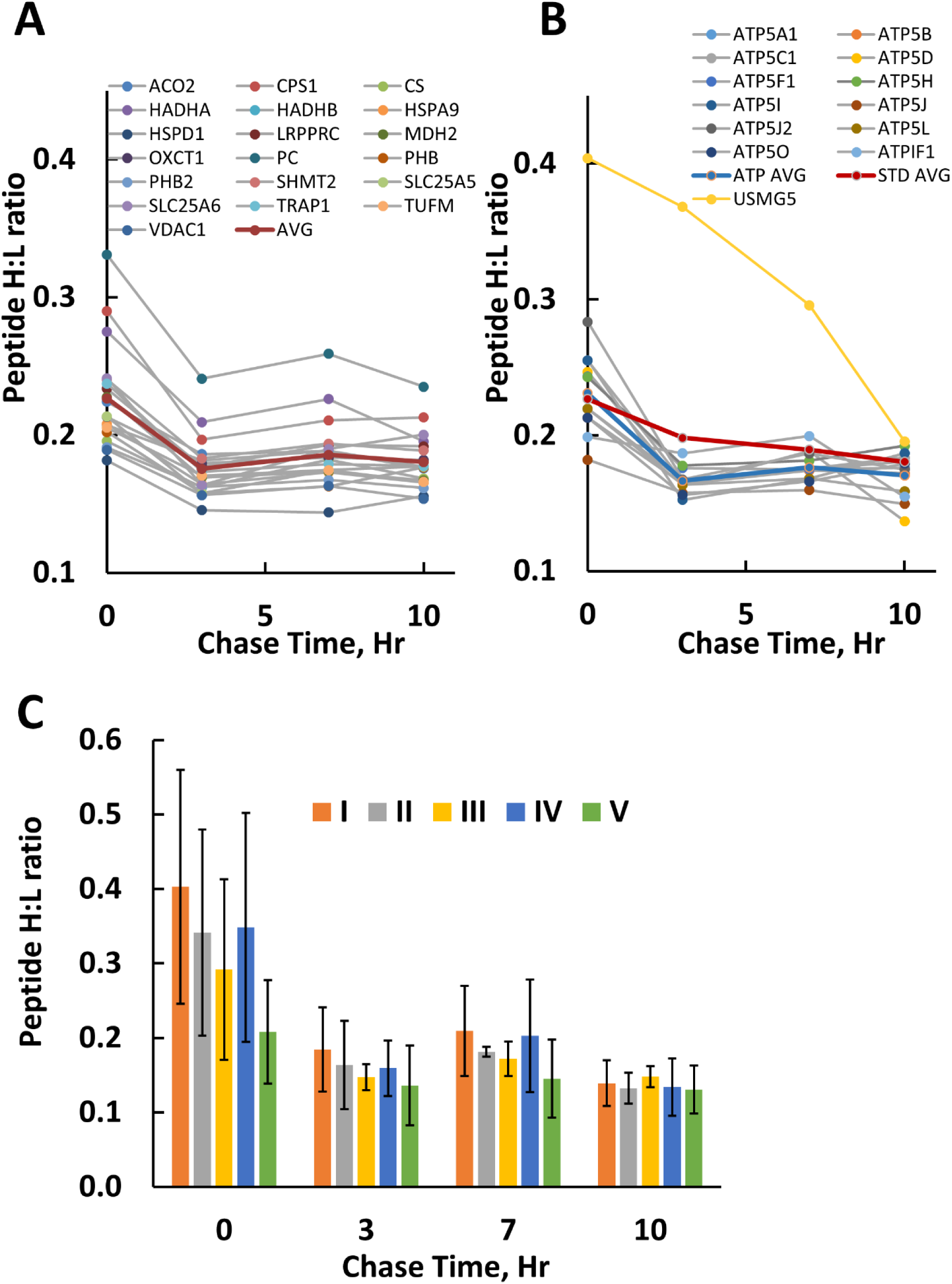
Protein H:L ratios in mitochondria decrease during chase intervals of 3, 7 and 10 h as excess over-synthesized protein is degraded. **A, B.** Peptide H:L ratios of standard proteins and RCV subunits, respectively, during the chase incubation. **C.** Comparison of the decreasing H:L ratios of peptides in subunits of each respiratory complex during the chase. Bars represent averages and error bars reflect standard deviation for the various groups of proteins. The initial large error bars reflect diversity in the levels of over-synthesis of individual proteins after pulse labeling while the smaller error bars after 10 h chase indicate that this variation is decreased as excess protein copies are degraded.

### Comparison of the rates of protein synthesis and turnover with RC assembly efficiency

To compare the kinetics of protein synthesis and import with those of RC assembly we used immunopurification (IP) to prepare RCI, RCIV and RCV after 6 hr SILAC labeling. We avoided use of shorter pulses for this purpose since we anticipated that the low extent of assembly would complicate detection of labeled nascent proteins using MS. Also, in preliminary experiments we tried using blue native PAGE gels for analysis of RC, but we considered that the limited loading capacity of these gels was not ideal for our goal of obtaining large numbers of peptide hits on as many proteins as possible.

IP proved to be a suitable method to analyze RC assembly as we observed peptide hits on the majority of structural subunits (Table S3; 39 of 44 RCI; 12 of 20 RCIV; 14 of 19 RCV). In initial experiments we found that commercial IP kits with antibodies conjugated to cross-linked agarose beads worked well for only RCV, not for RCI or RCIV in our hands. We did not include RCIII in this analysis. We found that a combination of immunopurification antibodies with Protein G magnetic beads gave superior results for RCI and RCIV as described in Experimental Procedures. As expected, hydrophobic membrane proteins such as COX3 and ATP5G as well as several membrane-domain RCI proteins were poorly represented in the proteomic analysis. Inferences related to assembly kinetics can be made with greater confidence when a protein is represented by a large number of peptide hits. In general, except as noted below, assembly factors were not observed in high yield in IP preparations.

In general we expect that the H:L ratios in assembled complexes should be lower than those in the total mitochondrial pool reflecting the time required for newly-synthesized imported protein copies to assemble into complexes capable of recovery by IP. We estimated that during a six-hour labeling period the total number of intact IP-competent RCI complexes should increase by about 19%, shown by the dashed horizontal lines in Fig. 3. If a complex requires a long interval for assembly we expect that the proteins that initiate assembly will have lower H:L ratios than those added late. This rationale is generally supported by our results.

**Figure 3.**
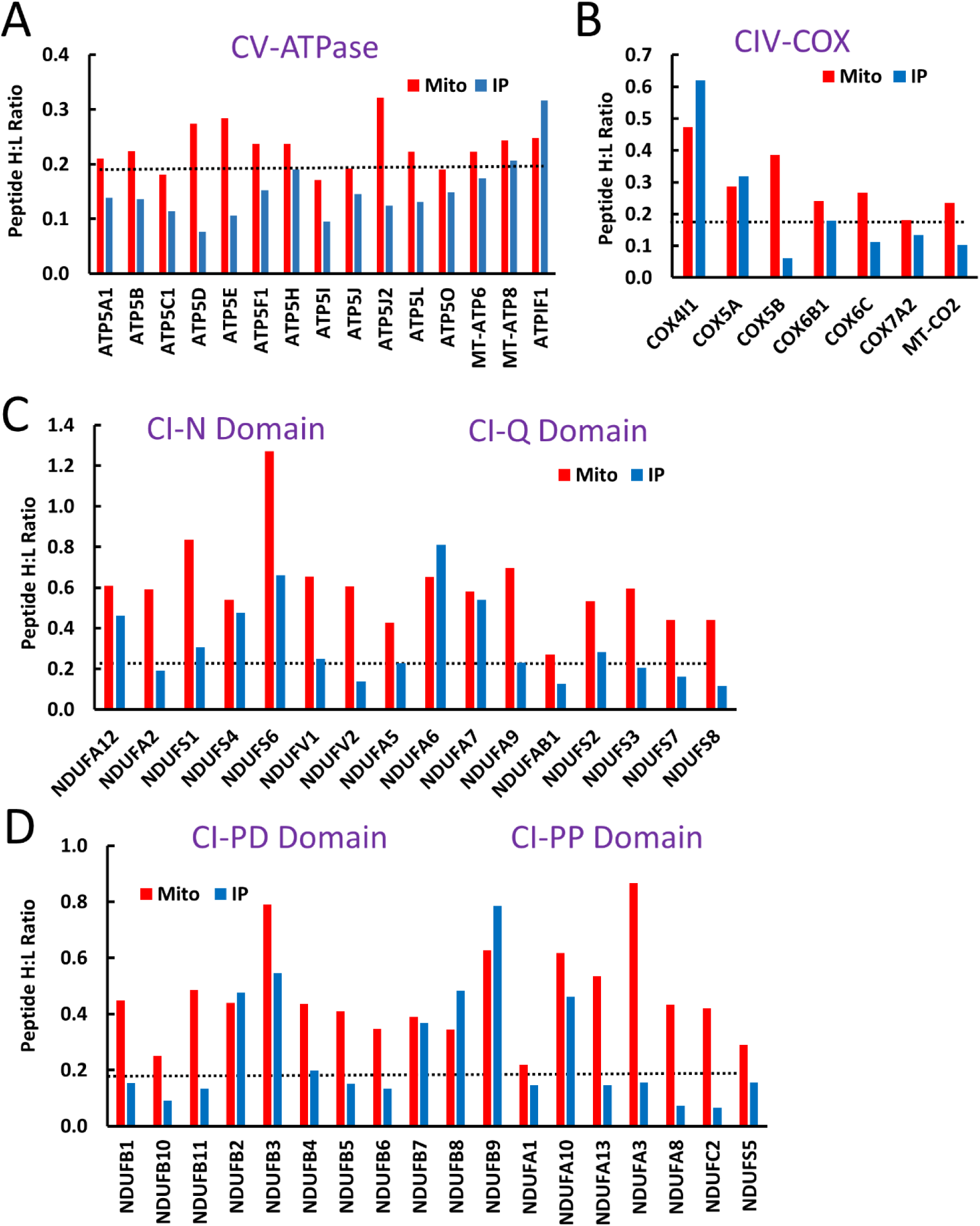
Comparison between the H:L ratios following 6 h pulse labeling in the total mitochondrial fraction (red bars) with those in immunoprecipitated respiratory complexes (blue bars). Note the change in scale on the Y-axis between panels reflecting much higher levels of synthesis and import of some proteins than others. The horizontal dotted lines indicate that approximately 19% of complexes are expected to be newly-synthesized during a 6 hr pulse interval. Error bars are not shown on these figures, but the primary data available in Table S3 indicates an average error of ±5% of the mean values for 45 RC proteins with good peptide coverage in both the total mitochondrial and IP samples. **A, B**. H:L ratios are shown for peptides in proteins identified in both total mitochondrial preparations (red) and samples immunoprecipitated with complex-specific antibodies (blue) in complex V and complex IV, respectively. **C, D.** H:L ratio results for respiratory complex I proteins similar to those in A and B for the matrix-exposed N and Q domains in C and the proximal (PP) and distal (PD) membrane domains in D.

### RCV

As anticipated from our simple pulse-chase studies, RCV is assembled relatively efficiently, as nascent peptides are quickly incorporated into immunoprecipitable complexes within the 6-hr labeling interval (Fig. 3A). The immunoprecipitated proteins with the greatest peptide coverage show H:L ratios 55 to 72% as high as their respective pulse-labeled total mitochondrial pools. The 8 proteins with the greatest peptide coverage in the IP samples had an average H:L ratio of 0.139±0.036, indicating a fairly uniform assembly rate. Although the mtDNA-encoded proteins MT-ATP6 and MT-ATP8 were represented by few peptides in both total mitochondrial preparations and IP samples, the H:L values observed for these peptides averaged 0.175 and 0.207, respectively, consistent with the behavior of other RCV subunits as well as the results of Lazarou et al. (5), who reported that nascent radioactively-pulse-labeled MT-ATP6 and 8 were rapidly incorporated into intact RCV. ATP5D and ATP5E showed the greatest discrepancy between the H:L ratios of the total mitochondrial pool and the assembled complexes, reflecting their slightly excessive rates of synthesis. The homogeneous rate of protein incorporation into IP-competent RCV reinforces the conclusion that RCV assembly is relatively rapid and efficient, as indicated by pulse-chase labeling results obtained for RCV proteins in the total mitochondrial fraction (Fig. 3A). The high content of newly-synthesized ATP5IF1 (ATPIF1) in the IP samples in Fig. 3A may reflect the fact that this is not a core subunit, but binds to inhibit the complex.

### RCIV

Comparison of the H:L ratios for immunoprecipitated RCIV proteins with the H:L ratios of these proteins in the total mitochondrial fraction in Fig. 3B presents a different pattern from that observed for RCV. RCIV assembly is initiated with the core mtDNA-encoded proteins MT-CO1, 2, and 3 (14). MT-CO1 and MT-CO3 were not observed in our IP experiments and are not included in Figure 3B. Greater peptide coverage was obtained for MT-CO2 revealing a total mitochondrial H:L ratio of 0.235 but a ratio of only 0.103 in IP samples. Similarly low H:L ratios were observed for some nuclear-encoded subunits such as COX5B and COX6. These low labeling indices suggest that these newly-synthesized proteins require a considerable time to progress to IP-competent complexes. In contrast, the H:L ratios observed for COX4IL and COX5B were essentially 100% of that in the total mitochondrial fraction. There are two possible explanations for this difference. One is that individual proteins may react with this antibody without bringing down the entire complex. According to the vendor, the epitope recognized by the IP antibody (ab109863) has not been mapped, but is considered to reside in one of these two proteins, or possibly at their interface. It is likely that this accounts for the high H:L ratios of COX4IL and COX5B peptides in our experiments. Based on this, we consider that the RCIV IP antibody is not an ideal reagent for this analysis. An alternative possibility is that COX41L, for example, may exchange readily into pre-existing RCIV assemblies. Previous studies suggest that early steps in RCIV assembly may occur at the inner boundary membrane juxtaposed to the outer membrane and that assembly may be completed as the complex migrates to the cristae membrane with COX4 added late (15, 16). Additional experiments may be required to determine whether nascent COX4IL and COX5B exchange into pre-existing RCIV complexes.

### RCI

RCI is the largest RC and has been studied extensively. RCI is built around a core of 14 proteins related to their E. coli counterparts, 7 membrane proteins encoded in mtDNA and 7 nucleus-encoded matrix-exposed proteins, surrounded by 30 so-called supernumerary subunits (8). In a broadly based genetic analysis, Stroud et al. (2017) found that five of these supernumerary proteins, NDUFV3, NDUFA7, NDUFA12, NDUFS4 and NDUFS6, are not strictly essential since knockout of these genes did not impair the ability of cells to grow using galactose as carbon source. Nevertheless, it should be noted that some of these may improve the stability or function of RCI, as suggested for a mouse NDUFS4 knockout (17). Another recent study (18) took advantage of the fact that incubation of cultured cells with chloramphenicol could deplete complexes that require subunits synthesized on mitoribosomes. Extensive protein complexome profiling during recovery of the electron transport chain after chloramphenicol washout established a detailed scheme for assembly of RCI through a series of sub-complexes. This study made the important observation that some nuclear-encoded proteins accumulate in these sub-complexes during the interval when mitochondrial protein synthesis is blocked. This is an issue that may be shared by numerous other studies that have monitored RC assembly following periods of treatment with translation inhibitors. Such studies have shown, for instance, that a chase interval as long as 24 hr is required for radioactively pulse-labeled mtDNA-encoded RCI subunits to appear in intact RCI (5). The time required to assemble RCI has not been well-studied without the use of protein synthesis inhibitors. We asked whether pulse SILAC would provide a complementary method to monitor RCI assembly without the confounding influence of perturbations using translation inhibitors. For the case of RCI the IP antibody used (ab109798) is known to recognize ND1 so that other subunits recovered are expected to reside in large complexes.

Figs. 3C and D compare the H:L ratios of SILAC pulse-labeled RCI subunits in IP complexes with the corresponding H:L ratios of the total mitochondrial pool of RCI. We assumed that if assembly is rather prolonged, many RCI subunits would contain only a low H:L ratio, since the subunits that seeded assembly of recently-completed complexes would probably have been synthesized before the pulse-labeling started. The majority of RCI subunits did indeed behave as expected. However, we noted several subunits with H:L ratios greater than or equal to twice that expected based purely on the growth rate. These include NDUFA12, NDUFS4, NDUFS6, NDUFA6 and NDUFA7 in the matrix-exposed N- and Q-domains. Several of these are included in the set of non-essential subunits noted above, and all of them were designated as late participants in RCI assembly by Guerrero-Castillo et al. (18). Thus, our data reinforces the conclusion that these proteins join the complex at a late stage in assembly. A notable difference is that treatment with chloramphenicol for several days in the experiments by Guerrero-Castillo et al. (18) most likely perturbed the pools of cytoplasmically synthesized subunits while pulse SILAC avoids this complication. We were surprised to find that some subunits in the membrane domain of RCI were also enriched in nascent polypeptides with high H:L ratio. These include NDUFA10, NDUFB2, NDUFB3, NDUFB7, NDUFB8, and NDUFB9. Fig. S2 shows the locations of these proteins as well as the late-binding matrix-domain proteins noted above within the RCI structure. NDUFA10 binds along with late-binding subunit NDUFA6 at the junction between the membrane and matrix-exposed domains. The other candidate late-binding membrane-domain subunits tend to associate with ND5 at the distal tip of the membrane domain. These proteins were also identified as late participants in RCI assembly by Guerrero-Castillo et al. (18). The high H:L ratios of these proteins in the IP samples after 6 hr of labeling is consistent with their late addition to the complex. Overall, our results obtained using simple SILAC labeling in unperturbed cells show considerable consistency with the elegant, but far more complex analysis of Guerrero-Castillo et al. (18), suggesting that this method has great potential to augment future studies of RCI assembly in wild-type and mutant cells.

### Nascent Rieske FeS protein and a module of proteins involved in FeS cluster biosynthesis co-IP with RCI

As noted above, Table S3 shows that accessory factors involved in RC assembly do not commonly co-IP with their respective complexes. However, the table identifies several counter-examples to this rule. NFS1 and several LYR-domain proteins co-precipitated with RCI, but not appreciably with other complexes. NFS1 and LYRM4 associate with NDUFAB1, which occurs in two copies in RCI and plays a dual role as acyl carrier protein (ACP) (7,19,20), although there is a free pool of ACP in mitochondria as well (21). Interestingly, after 6 hr SILAC labeling the H:L ratio of NDUFAB1 (ACP) in immunoprecipitated RCI is lower than that of the two LYR-domain proteins NDUFA6 and NDUFB9 (Fig. 3). This may indicate that newly-synthesized NDUFAB1 mixes with the pool of pre-existing ACP prior to incorporation in RCI. The NFS1-LYRM4-NDUFAB1 complex also associates with other proteins involved in RCI assembly including LYRM1, LYRM2, LYRM7 and two factors implicated in RCV assembly, ATPAF2 and FMC1 (C7orf55) (22, 23). All of these were represented by numerous peptides in our RCI IP samples (boxed in purple in Table S3). Whether this association of FeS synthesis machinery with RCI is involved in introduction of the numerous FeS complexes into RCI subunits is not well known. Genetic knockout of the *Y. lipolytica* NDUFA6 was found to impair ubiquinone metabolism but not to dramatically affect Fe content of RCI (24). Two groups have recently reported structures of complexes in which human NFS1-LYRM4, and in one case, ISCU as well, adventitiously associated with E. coli ACP (19, 20). Although these structures were not in agreement, both of them as well as the human RCI structure show that the human or E. coli ACP consistently associates through the respective LYR protein using a similar interaction interface, as recently reviewed (25). Thus, ACP cannot interact simultaneously with LYRM4 (ISD11) in the FeS complex and also with either LYRM3 (NDUSB9) and/or LYRM6 (NDUFA6) in the mature RCI complex. Whether dynamic interactions are possible involving RCI assembly intermediates will require additional investigation to understand the structural basis of the interaction between RCI components and FeS cluster proteins.

We were also surprised to find that the RCI IP also revealed a significant number of peptide hits for UQCRFS1, the Rieske FeS protein from RCIII, and, remarkably, that this RCI-associated fraction was highly enriched in newly-synthesized protein. The H:L ratio of this UQCRFS1 associated with RCI was over twice as great as the total mitochondrial pool of UQCRFS1 (Fig. 4A). UQCRFS1 is known to be imported into the matrix for processing, including addition of the FeS cluster, before it is exported to the intermembrane space with the assistance of the AAA-ATPase BCS1L to be inserted into a late RCIII assembly intermediate (26). Protein interaction analysis using the STRING database (https://string-db.org/) indicates that UQCRFS1 shares extensive interactions with the LYR domain and FeS complex proteins noted above (Figure 4B). It is tempting to suggest that the FeS complexes associated with RCI also participate in introduction of FeS complexes in newly-synthesized UQCRFS1. However, it should be noted that the full complement of proteins required for synthesis and incorporation of FeS clusters is not represented in our IP samples (27, 28). A more extensive investigation of the association of this FeS-assembly complex and of UQCRFS1 with RCI is beyond the scope of our current investigation. These interactions may contribute to the known crosstalk between RCI and RCIII assembly (29).

**Fig. 4.**
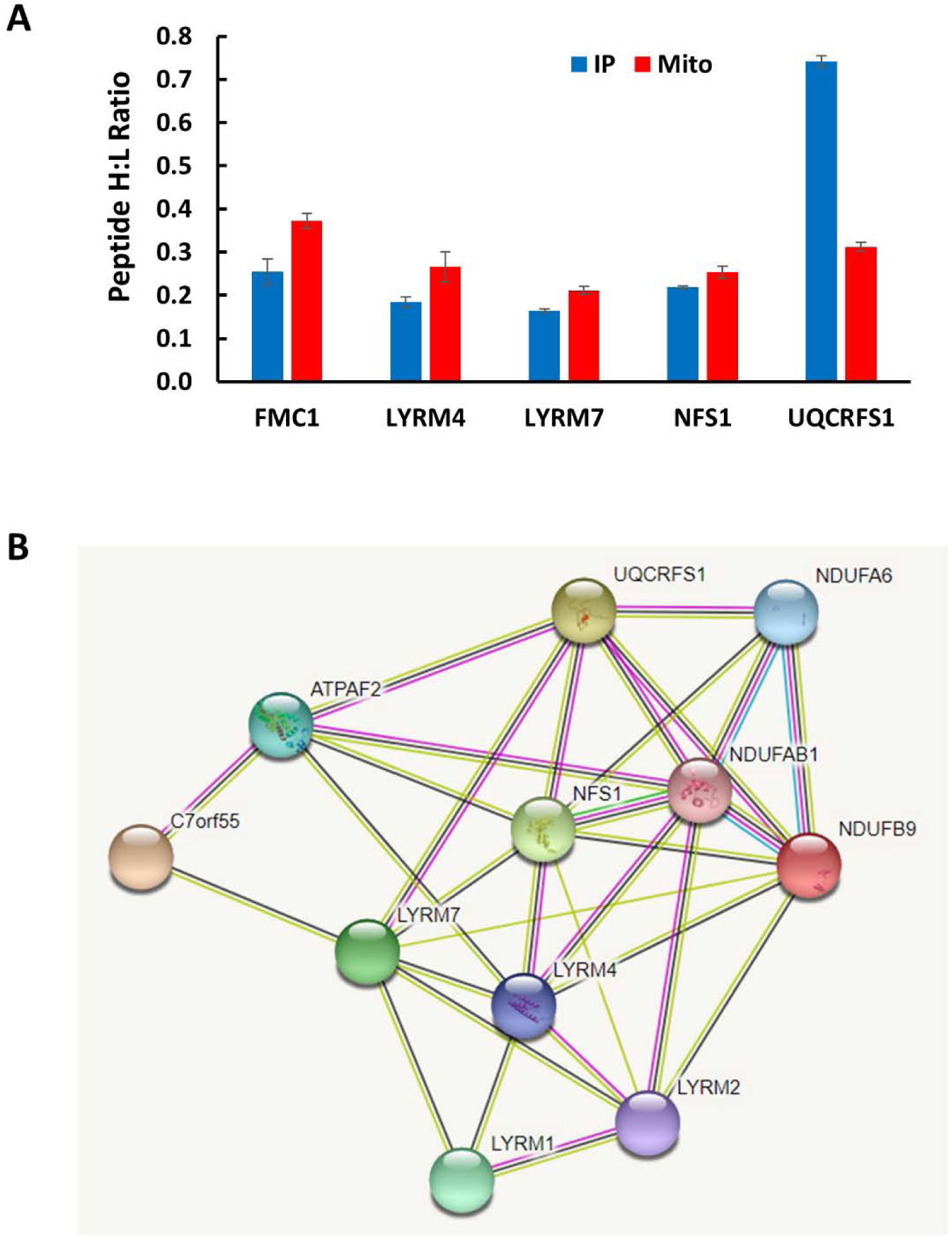
Newly-synthesized UQCRFS1 and a module of LYR-domain proteins are selectively immunoprecipitated with RCI. **A.** IP of RCI from lysates of mitochondria from cells labeled for 6 hr recovered several proteins that are not structural subunits of RCI (Table S3). For proteins represented by statistically significant numbers of peptides (>20) the chart shows the mean values (+/-standard error) for the total mitochondrial pool (red) and for the fraction of protein recovered by RCI IP (blue). UQCRFS1 stands out as a member of another RC, RCIII, with a significant over-representation of newly-synthesized protein. **B.** The network of protein interactions among NDUFAB1, its associated LYR domain proteins in RCI (NDUFA6 and NDUFB9), UQCRSF1, NFS1 and other assembly factors recovered in RCI IP samples visualized using STRING (https://string-db.org).

## Conclusions

The structures of each of the major complexes in the mammalian respiratory chain are now clearly defined, but our understanding of the assembly of these macromolecular machines remains incomplete. Assembly requires independent synthesis of nearly 70 cytoplasmically-translated proteins that must be co-assembled with 13 mtDNA-encoded subunits in a coordinated manner. Excessive synthesis of subunits required in strict stoichiometry must lead to either an accumulation of unassembled subunits or to their turnover. While we are beginning to see how regulated translation of nuclear-encoded subunits may help to coordinate assembly (16,30,31), understanding of these processes is incomplete.

We employed pulse-chase SILAC experiments to test whether this approach can complement other existing methods to study the dynamics of respiratory complex assembly. The SILAC method avoids the use of radioisotopes, does not rely on use of protein synthesis inhibitors that may result in imbalance in the steady-state pools of protein subunits and yields information on large numbers of mitochondrial proteins in each experiment. We find that proteins synthesized and imported at higher rates than required to keep pace with cell growth also turn over more rapidly.

We find remarkable differences in the rates at which individual components of respiratory complexes are synthesized and imported into mitochondria. This is not unexpected since we recently reported that assembly of the mitochondrial ribosomes involves a degree of over-synthesis of some of the 80 nucleus-encoded proteins (2). We reasoned that this excess synthesis might be useful to provide an adequate supply of parts for the assembly process. This rationale appears to apply to RC complex assembly as well, but the over-synthesis of ribosomal proteins is quite modest compared to some of the respiratory complex subunits. The rates of synthesis of NDUFS6 and the complex IV protein NDUFA4 are at least 5 times as rapid as required to sustain cell growth, so that about 80% of the newly-synthesized copies of these proteins are fairly quickly degraded (Table S2). On average all of the subunits of the matrix-exposed domains of RCI are synthesized two to three times as rapidly as required. In contrast, all of the core subunits of RCV are synthesized at rates only slightly in excess of growth demands.

RCI contains more than twice as many subunits as any other RC, such that its assembly is far more complex than that of RCV (18, 32). Our experiments establish that pulse-chase SILAC provides a relatively simple method to investigate this assembly process. Moreover, it provides an opportunity to track the fate of newly-synthesized proteins during the construction process. This has revealed the considerable over-synthesis of many of the supernumerary subunits of RCI and the unanticipated association of a network of LYR-domain proteins and FeS synthesis participants with RCI and/or its assembly intermediates. It is particularly noteworthy that we discovered a transient association of newly-synthesized copies of the Rieske FeS protein with RCI. Future experiments beyond the scope of our current work may probe these interactions in greater detail. It will be particularly interesting to apply these methods to study complex assembly in cells derived from patients with mutations in RC complex proteins or assembly factors.

## Experimental Procedures

### Cell growth and SILAC labeling

HeLa cells were grown in monolayer culture in DMEM (ThermoFisher Scientific) supplemented with 10% fetal bovine serum (Atlanta Biologicals), 2.5 μg/ml amphotericin B (Corning), 1% penicillin/streptomycin (ThermoFisher Scientific). SILAC labeling used kits from ThermoFisher Scientific with DMEM lacking lysine and arginine supplemented with 30 mg/L ^13^C6-arginine and 50 mg/L ^13^C6-lysine with 10% dialyzed fetal bovine serum. Labeling was initiated after one rinse of tissue culture plates with pre-warmed phosphate buffered saline and 2 rinses with Hanks’ balanced salt solution before addition of ^13^C-DMEM. Chase incubation with normal supplemented DMEM was similarly preceded with wash steps. Medium changes for SILAC pulse and chase incubations began at a time judged to permit conclusion of incubations before cells reached 90% confluence.

### Mitochondrial purification and RC immunoprecipitation (IP)

Mitochondria were purified as described (2). For IP of RC, 100 to 400 μg mitochondria were lysed on ice for 30 min with 2.5% n-dodecyl-β,D-maltoside in 400 or 600 μl buffer A20 containing 20 mM NaCl, 50 mM imidazole-HCl pH 7, 2 mM 6-aminocaproic acid, 1 mM EDTA and HALT protease inhibitor (ThermoFisher Scientific). Lysates were clarified by centrifugation at 15,000g for 20 min at 4 ⁰C. The lysate was supplemented with 10 to 12 μg antibodies against RCI (Abcam ab109798) or RCIV (Abcam ab109863) or with 15 μl of immobilized antibodies against RCV (Abcam ab109715). For RCI or RCIV IP, 60 to 120 μl of Protein G Dynabeads (Thermo Fisher 10003D) washed with PBS was added after 1 hr incubation of proteins with free antibodies and incubation was continued for at least 1 additional hour. In some cases, RCI or RCIV IP using Protein G magnetic beads was combined with RCV IP using crosslinked agarose beads which allowed the magnetic beads to be separated on a magnet before the agarose beads were recovered by centrifugation at 3000g for 1 min. Beads were washed with PBS containing 0.05% n-dodecyl-β,D-maltoside three times and proteins were eluted with successive washes with 50 μl 0.2 M glycine, pH 2.5. Proteins were concentrated using chloroform-methanol precipitation (33) and processed for proteomic analysis.

### Protein analysis by LC-MS/MS

Proteins were fragmented with trypsin and peptides were detected by LC-MS/MS on an AB-Sciex 5600Plus orthogonol quadrupole TOF mass spectrometer as described (2, 34). Peptides were sorted to eliminate those with low heavy or light peptide signals, confidence scores <90% and those that might represent another peptide. Peptides with inappropriately high H:L ratios in the highest 1% of the distribution were manually removed as outliers. Peptide information was imported into SAS JMP 13 software to generate average H:L ratios and standard error calculations using robust fitting that calculates geometric means for data distributions (35). The mathematical model relating the expected H:L ratio of a protein (P) to the labeling time (t) and growth rate (Tg) was developed in published supplementary material (2), yielding the relation P^H^/L = (e^ln2*t/Tg^ – P_L_(0)e^ln2*t/t1/2^)/e^ln2*t/t1/2^ that relates the H:L ratio for a protein (P) to the duration of the labeling time (t) and the cell generation time (Tg).

## Acknowledgements

I thank Anne Ostermeyer-Fay for technical assistance and Miguel Garcia-Diaz and Sandeep Mallipattu for comments on the manuscript. This work was supported by NIH grant 1R01 GM112790 and by a research grant from the United Mitochondrial Disease Foundation.

## Supporting information

1. Tables of all pulse label peptides with summary
2. Tables of al chase peptides with summary
3. Tables of all RC IP peptides

## Supplemental Figures

**Fig. S1.**
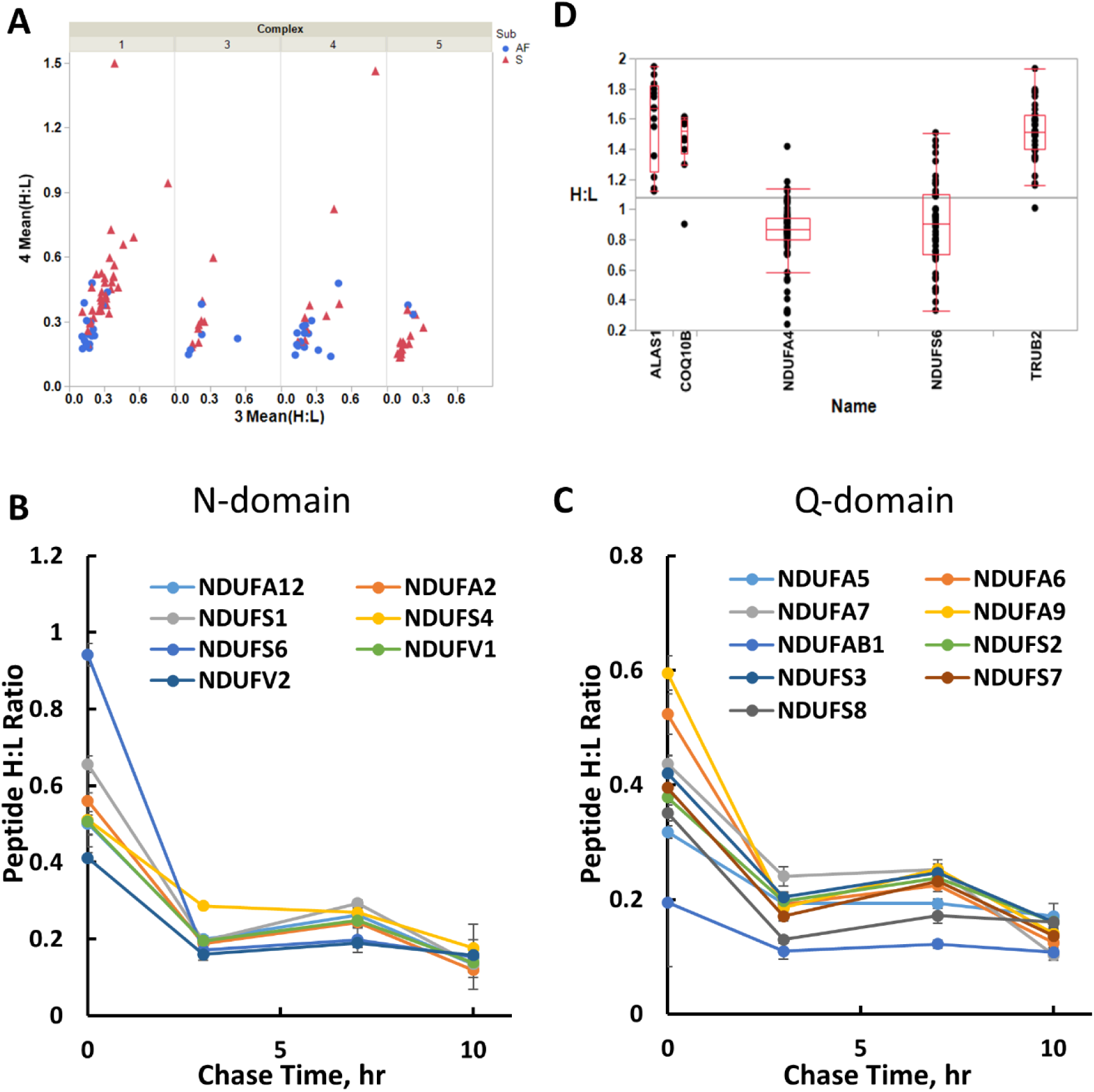
Some proteins are synthesized and imported at particularly high rates and can be turned over with biphasic kinetics. **A**. Scatterplots of the H:L ratios of individual proteins in RCI, RC3, RC4 and RC5 after SILAC pulse labeling for 3 hr (abscissa) or 4 hr (ordinate). Complex structural subunits (red) generally have higher H:L ratios than accessory factors (blue). **B, C.** Time courses for the decline in H:L ratio during the chase following a 4 hr SILAC labeling for RCI subunits in the N- and Q-domains, respectively. **D.** Examples of the distribution of H:L ratios of peptides observed following 3 hr pulse labeling for the indicated highly synthesized proteins. Boxplots show median and quantile values for peptides from individual proteins.

**Fig. S2.**
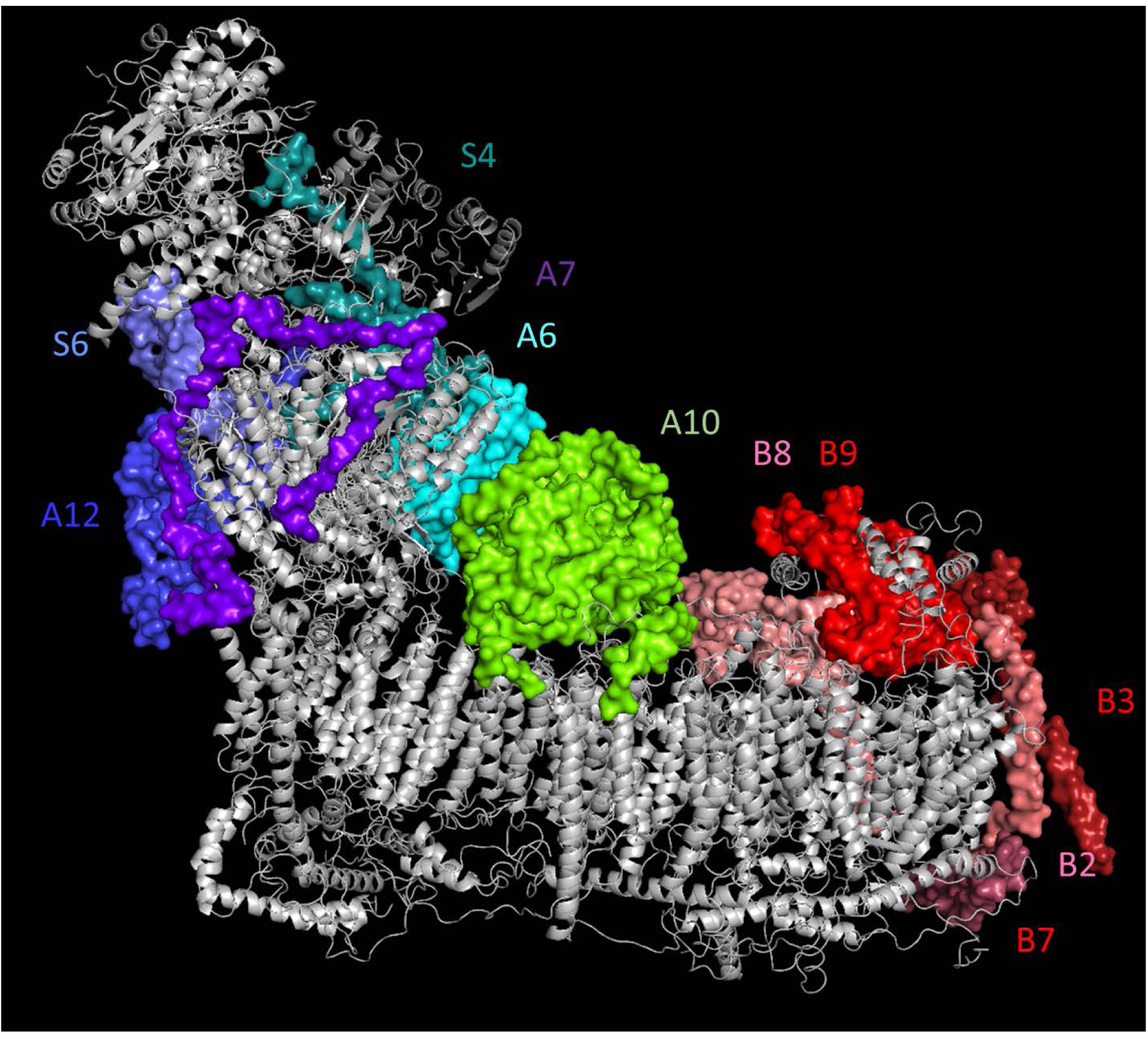
Structure of RCI showing the locations and identities of subunits with the highest H:L ratios (greatest amount of newly-synthesized protein) in immunoprecipitated RCI complexes after 6 hr pulse labeling. Subunits added late within the N and Q domains are shown with surface coloration in shades of blue. Subunits highly labeled in the PD domain are shown with surface coloration in shades of red. NDUFA10 (green) is shown as the only highly labeled PP-domain protein. Names of RCI subunits are truncated for clarity. Note that NDUFA6 and NDUFB9 are the LYR-domain proteins LYRM6 and LYRM3, respectively, in contact with the two subunits of NDUFAB1 (ACP) as discussed in the text.

